# Regulation of *yefM/yoeB* toxin antitoxin system is independent of ppGpp and inorganic polyphosphate in *Escherichia coli*

**DOI:** 10.1101/021162

**Authors:** Bhaskar Chandra Mohan Ramisetty

## Abstract

Bacterial persistence is a phenomenon wherein small proportion of a bacterial population attains transient antibiotic tolerance likely by virtue of metabolic minimization. Type II Toxin–Antitoxin systems (TAs), small overlapping bicistronic negative auto-regulons, were recently shown to induce the persistence state. Maisonneuve et al., 2013 reported that TAs are activated by a regulatory cascade consisting of stochastic accumulation of ppGpp leading to accumulation of inorganic polyphosphate (polyP). PolyP supposedly is essential for Lon protease-dependent-degradation of antitoxins resulting in activation of toxins and induction of persistence phenotype. In contrast, using semi-quantitative primer extension, we show that transcriptional up-regulation of *yefM/yoeB* loci, one of the well characterized TAs of *Escherichia coli*, is independent of ppGpp and polyP. Similarly, we show that chromosome-encoded YoeB-dependent target mRNA cleavage is independent of polyP. Our results and meta-analysis of literature we conclude that the regulation of *yefM/yoeB* TAs is independent of ppGpp and polyP.

## INTRODUCTION

Toxin–antitoxin systems (TAs) are operons consisting of two or three adjacent genes which code for a toxin, which has the potential to inhibit one or more cellular processes, and an antitoxin. The antitoxin forms a complex with the toxin and suppresses the lethality of the toxin. Prokaryotic DNA sequence database mining showed that TAs are abundant in bacterial and archaeal chromosomes often in surprisingly high numbers (Anantharaman and Aravind 2003, Pandey and Gerdes 2005, Shao, et al. 2011). Based on the gene products, either RNA or protein, TAs are divided into 5 types (Goeders and Van Melderen 2014) of which Type II are the most predominant and well characterized. Type II TAs encode two proteins referred to as toxin and antitoxin. They are the most predominantly encoded type by bacterial genomes and plasmids. The toxin has the potential to inactivate vital cellular targets while the antitoxin has the potential to sequester toxins off the cellular targets by forming a toxin-antitoxin complex. Toxins and antitoxins also have the autoregulatory function wherein the TA complex binds to the operator present upstream of the TA operon and results in repression. The antitoxin is highly unstable and its relative concentration plays a critical role in transcriptional autoregulation as well as regulation of toxin activity. The decrease in antitoxin concentration is a prerequisite for transcriptional activation of TAs. The significance of TAs multiplicity on prokaryotic genomes and their physiological role is highly debated. Many plasmids also encode TAs whose gene products have the ability to inhibit the growth of the cells cured of TA-encoding plasmids and thereby increase the population of plasmid-containing cells (Gerdes, et al. 1986). Chromosomal TAs were discovered in studies dealing with stringent response and persistence. Stringent response, a response elicited in cells under amino acid starvation, is characterized by accumulation of ppGpp alarmone catalyzed by RelA upon stimulation by uncharged tRNA at the ribosomal A site (Cashel, et al. 1996, Haseltine and Block 1973, Lund and Kjeldgaard 1972, Wendrich, et al. 2002). Accumulation of ppGpp modulates RNA polymerase resulting in reduction of rRNA synthesis and thus prevents frivolous anabolism (Artsimovitch, et al. 2004, Barker, et al. 2001). Several mutants deficient/altered in stringent response were shown to be mutants of *relBE*, a TAs encoding an antitoxin (RelB) and a ribosome dependent endoribonuclease toxin (RelE) (Christensen, et al. 2001, Gotfredsen and Gerdes 1998). Persistence, a phenomenon of non-inheritable antibiotic tolerance, is the second instance in which genes belonging to the TA family were recognized. Some mutants, high persister mutants (*hip*), of *Escherichia coli* formed more number of persisters than the wild type. These *hip* mutations mapped to the *hipA* locus (Moyed and Bertrand 1983) which is now recognized as a genuine TAs encoding HipA toxin and HipB antitoxin (Germain, et al. 2013, Kaspy, et al. 2013, Korch, et al. 2003). A recent study shows an attractive link between TAs, stringent response and persistence; ppGpp, through inorganic polyphosphate (polyP), activates TAs resulting in induction of persistence (Maisonneuve, et al. 2013).

The crucial link between ppGpp and TA-mediated persistence is the essentiality of polyP for the degradation of antitoxins. During stringent response, polyP accumulates due to ppGpp-mediated inhibition of exopolyphosphatase (PpX) (Kuroda, et al. 1997). The presence or absence of polyP determines the substrate specificity of Lon protease (Kuroda, et al. 2001). Maisonneuve et al., 2013 have shown that polyP is essential for Lon-dependent degradation of YefM and RelB antitoxins resulting in increased persistence. YefM is the antitoxin encoded by *yefM/yoeB* TAs, a well-characterized Type II TAs. YoeB, the toxin, is a ribosome-dependent endoribonuclease (Christensen-Dalsgaard and Gerdes 2008, Feng, et al. 2013) that cleaves mRNA. YefM forms a complex with YoeB resulting in inhibition of endoribonuclease activity of YoeB (Cherny, et al. 2005, Kamada and Hanaoka 2005) and also in mediating transcriptional autorepression (Kedzierska, et al. 2007). However, earlier studies indicate that transcriptional regulation of TAs, like *relBE* and *mazEF* systems, is independent of ppGpp (Christensen, et al. 2001, Christensen, et al. 2003) and likely of polyP as well. Hence, in this study we analyzed the essentiality of polyP in degradation of YefM antitoxin by studying the promoter activity of *yefM/yoeB* loci and endoribonuclease activity of chromosomally encoded YoeB.

**Figure 1.**
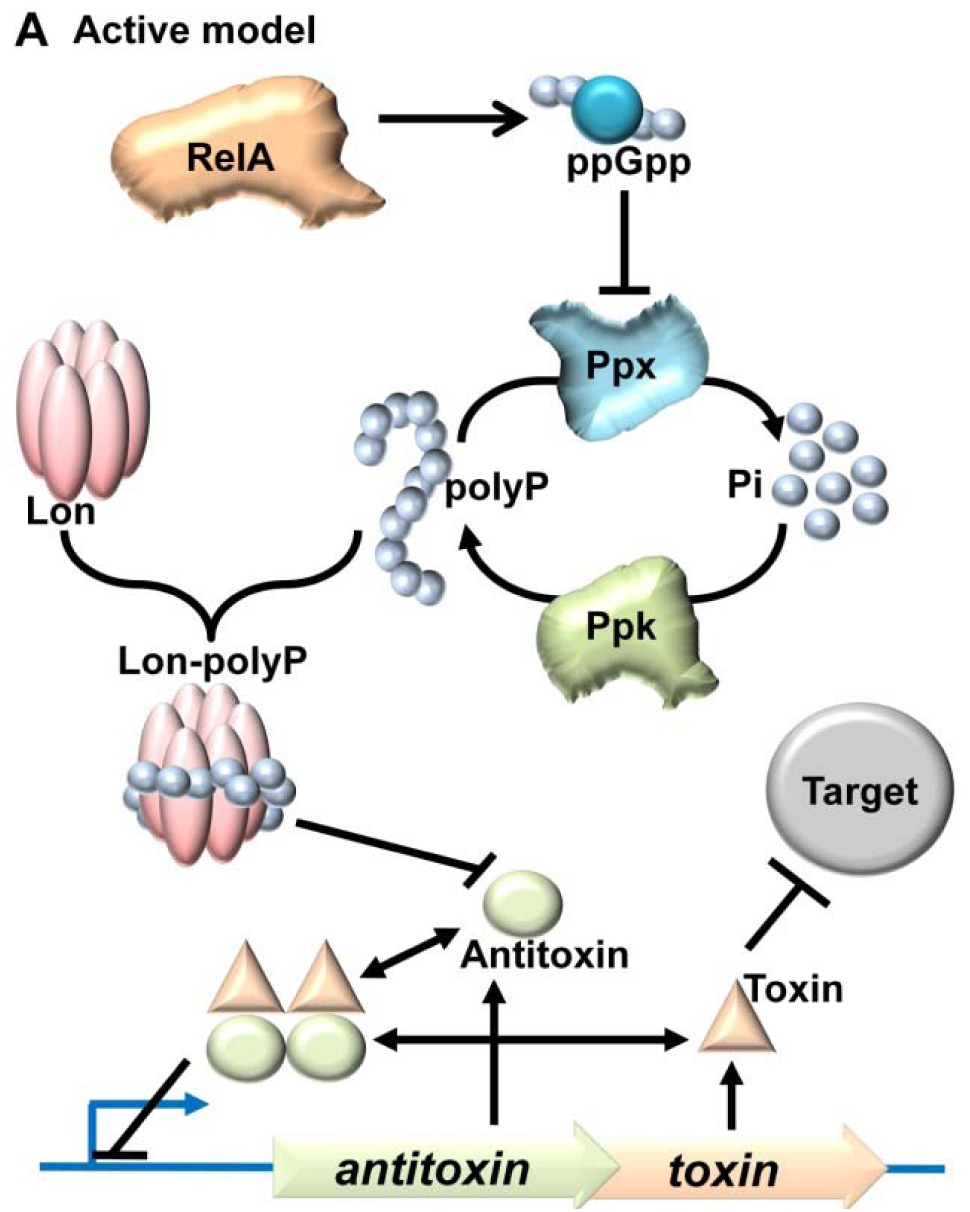
Active model of TAs regulation. RelA, when activated stochastically or during amino acid starvation, synthesizes the ppGpp. Accumulation of ppGpp inhibits the degradation of polyP into inorganic phosphate. Hence, due to continual synthesis by Ppk, polyP accumulates in the cell. PolyP then modulates the substrate specificity of Lon protease, specifically targeting antitoxin proteins for degradation. This is hypothesized to render the toxin free to act on its target and confer persistence.

Complex regulatory mechanisms for TAs activation and numerous physiological roles for chromosomal TAs were proposed by different groups. The most recent report is that TAs are involved in persistence and that ten different TAs are regulated by ppGpp via polyP modulated Lon dependent cleavage of antitoxins {Maisonneuve, 2013 #8157}. The autoregulatory bicistronic circuitry of Type II TA loci is primarily dependent on the concentration of antitoxin as it regulates the transcription and also the phenotypic manifestation of the toxins. During stress conditions the antitoxin levels reduce which could either be due to decrease in the production of antitoxin and/or increase in the degradation of antitoxin. ‘Passive’ and ‘Active’ models were described to explain the possible mechanisms in the reduction of antitoxin concentration (Gerdes, et al. 2005). In the “passive model,” the reduction in antitoxin concentration is due to the inhibition of translation while in the “active model” it is due to enhanced proteolysis of antitoxin. In the active model the antitoxins are hypothesized to be specifically targeted for degradation by its cognate protease. It was speculated that polyP has a role in the active regulation of TAs, by acting as a Lon stimulant to specifically degrade antitoxins (Gerdes, et al. 2005). In a recent report it was shown that *yefM/yoeB* and *relBE* systems are under the “active” control of ppGpp through polyP (Maisonneuve, et al. 2013).

## RESULTS

The transcriptional upregulation of *yefM/yoeB* loci, or any typical TAs, is inversely proportional to the relative concentration of YefM. This is because TA proteins autoregulate their promoter/operator; at higher antitoxin concentration the promoter repression is more and vice verse. Hence any transcriptional activation from *yefM/yoeB* operon indicates a decrease in antitoxin concentration which could be a result of either increased proteolysis or decreased translation of YefM. Therefore, quantification of the TA mRNA is a good indicator of antitoxin concentration in the cell. To test the essentiality of polyP in Lon-dependent degradation of YefM in vivo, we employed semi-quantitative primer extension (Christensen, et al. 2001, Christensen, et al. 2003) of YefM mRNA. This assay has the advantage of a holistic transcriptional regulatory scenario of TAs without employing any genetic manipulations within the TA circuitry, thus avoiding artifacts.

### polyP is not required for upregulation of *yefM/yoeB* loci during amino acid starvation

To test the role of ppGpp and polyP in the regulation of *yefM/yoeB* system, we performed amino acid starvation experiments using serine hydroxymate (SHX) and analyzed the transcription of *yefM/yoeB* loci using semi-quantitative primer extension using a YefM mRNA-specific primer. Exponentially growing *E. coli* strains MG1655 (Wild type), Δ*lon*, Δ*ppk*Δ*ppx* and Δ*relA*Δ*spoT*, were treated with 1 mg/mL of SHX to induce serine starvation. Δ*ppk*Δ*ppx* and Δ*relA*Δ*spoT* strains are deficient in accumulating polyP and ppGpp, respectively (Crooke, et al. 1994, Xiao, et al. 1991). In the wild type (WT) strain, we found a dramatic increase (16 fold) in the transcription of *yefM/yoeB* loci while in Δ*lon* strain there was no change (Figure 2A). Interestingly and importantly, we found an increase in transcription of *yefM/yoeB* loci in Δ*ppk*Δ*ppx* as well as in Δ*relA*Δ*spoT* strains, similar to that of the WT. This observation indicates that *yefM/yoeB* transcriptional control through YefM is under the control of Lon and is independent of polyP and ppGpp. In fact, earlier studies reported ppGpp-independent transcriptional upregulation of *relBE* (Christensen, et al. 2001) and *mazEF* systems (Christensen, et al. 2003) during SHX-induced starvation experiments.

**Figure 2.**
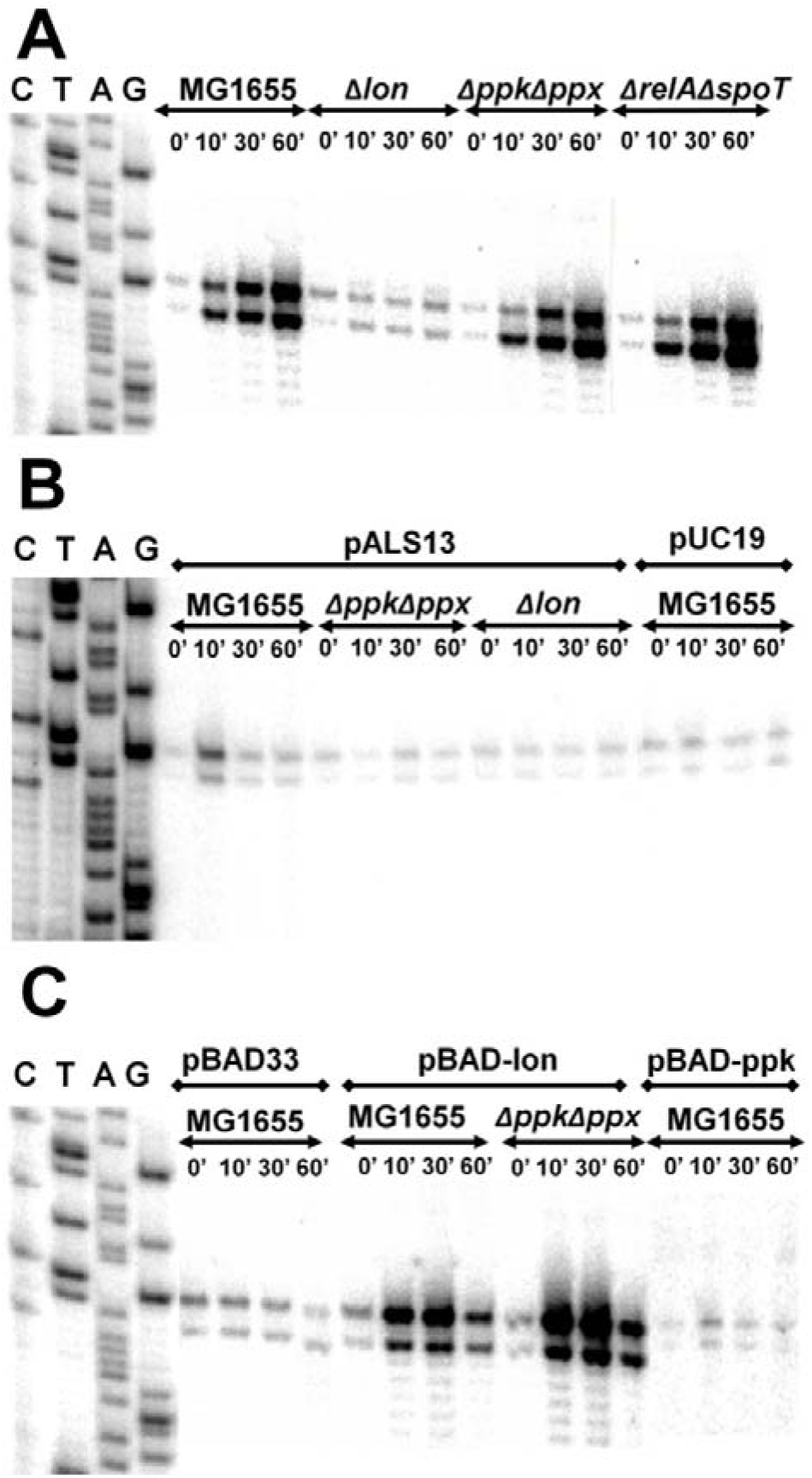
Transcriptional upregulation of *yefM/yoeB* loci is independent of polyP: (**A**) Exponentially growing (0.45 of OD_450_) cultures of MG1655, Δ*lon*, Δ*ppkppx* and Δ*relA*Δ*spoT* were treated with 1 mg/ml of serine hydroxymate. Total RNA was isolated at 0, 10, 30 and 60 minutes and semi-quantitative primer extension was performed using YefM mRNA-specific primer (YefMPE-2). (**B**) pALS13 plasmid, containing truncated *relA* gene downstream of the *lac* promoter, was transformed into MG1655, Δ*lon* and Δ*ppkppx* strains. All the strains were grown to exponential phase and *relA’* was overexpressed using 1 mM Isopropyl β-D-1-thiogalactopyranoside (IPTG). Total RNA was purified from samples taken at 0, 10, 30 and 60 minutes after addition of IPTG. Primer extension was performed as in (A). (**C**) MG1655 strain was transformed with pBAD33 or pBAD-*lon* or pBAD-*ppk* plasmids and Δ*ppkppx* strain was transformed with pBAD-*lon*. Overnight cultures were diluted and grown to 0.45 OD_450_ in LB medium supplemented with glycerol as carbon source at 37 °C. 0.2% arabinose was added to induce overepxression of *lon* or *ppk*. Samples were collected at different intervals and semi-quantitative primer extension performed as in (B).

### Lon protease-induced transcriptional upregulation of *yefM/yoeB* is independent of polyP

To further analyze the role of ppGpp and polyP in *yefM/yoeB* regulation we used ectopic overexpression of *relA*′ (Svitil, et al. 1993), which encodes a truncated RelA capable of ribosome-independent synthesis of ppGpp, in different strains. WT, Δ*ppk*Δ*ppx* and Δ*lon* strains, transformed with pALS13 (Svitil, et al. 1993), were grown to mid log phase and the expression of *relA*′ was induced by the addition of IPTG. In our primer extension analysis of *yefM/yoeB* trasncription, we found that there was transient increase (7 fold) in the transcription during the first 10 minutes but decreased back to basal levels. However, in Δ*ppk*Δ*ppx* and Δ*lon* strains, there was no change over a time period of 60 minutes (Figure 2B). This indicates that there is a polyP-dependent transient increase in *yefM/yoeB* transcription upon overproduction of ppGpp during the exponential growth phase. We also carried out overexpression of Lon protease in MG1655 and Δ*ppk*Δ*ppx* strains to check if polyP has a role in the regulation of *yefM/yoeB* system and found that transcription of *yefM/yoeB* increased similarly in both MG1655 and Δ*ppk*Δ*ppx* strains (Figure 2C). Similarly, upon ectopic overexpression of *ppk* we found a transient and marginal increase in transcription from *yefM/yoeB* loci in samples taken at 10 minutes but reduced back to basal levels by 30 minutes.

### YoeB-mediated cleavage of mRNA upon overexpression of Lon is independent of polyP

Transcriptional upregulation of *yefM/yoeB* operon does not necessarily mean that YoeB is free to cleave its target mRNA. To date, chromosomal YoeB-dependent mRNA cleavage has been observed only upon ectopic overproduction of Lon protease (Christensen, et al. 2004). The ectopic overexpression of Lon degrades YefM, leaving YoeB free to manifest its endoribonuclease activity. Since it was shown that Lon-mediated degradation of YefM is dependent on polyP (Maisonneuve, et al. 2013), it is interesting to see if polyP is essential to render YoeB free by promoting the degradation of YefM. First, we overexpressed Lon protease in WT, Δ*ppk*Δ*ppx*, Δ*yefM/yoeB* (MG1655 derivate with *yefM/yoeB* deletion) and Δ*5* (MG1655 derivate in which 5 TAs are deleted) strains and mapped for cleavage sites in Lpp mRNA by primer extension as reported in earlier studies (Christensen, et al. 2004). We found that Lpp mRNA is cleaved at the second codon of AAA site in WT and Δ*ppk*Δ*ppx* strains but not in Δ*yefM/yoeB* and Δ*5* strains (Figure 3). As evident from Figure 2C, there was a transient upregulation of *yefM/yoeB* transcription upon overexpression of *ppk* and *relA*. Hence, we overexpressed *ppk* in exponentially growing cultures of WT, Δ*lon*, Δ*yefM/yoeB* and Δ*5* strains. We could not detect any YoeB-dependent cleavage of Lpp mRNA upon ectopic overexpression of *ppk* in any of the strains. These results indicate that polyP is not required to render YoeB free of YefM to manifest YoeB dependent cleavage during Lon overproduction.

**Figure 3.**
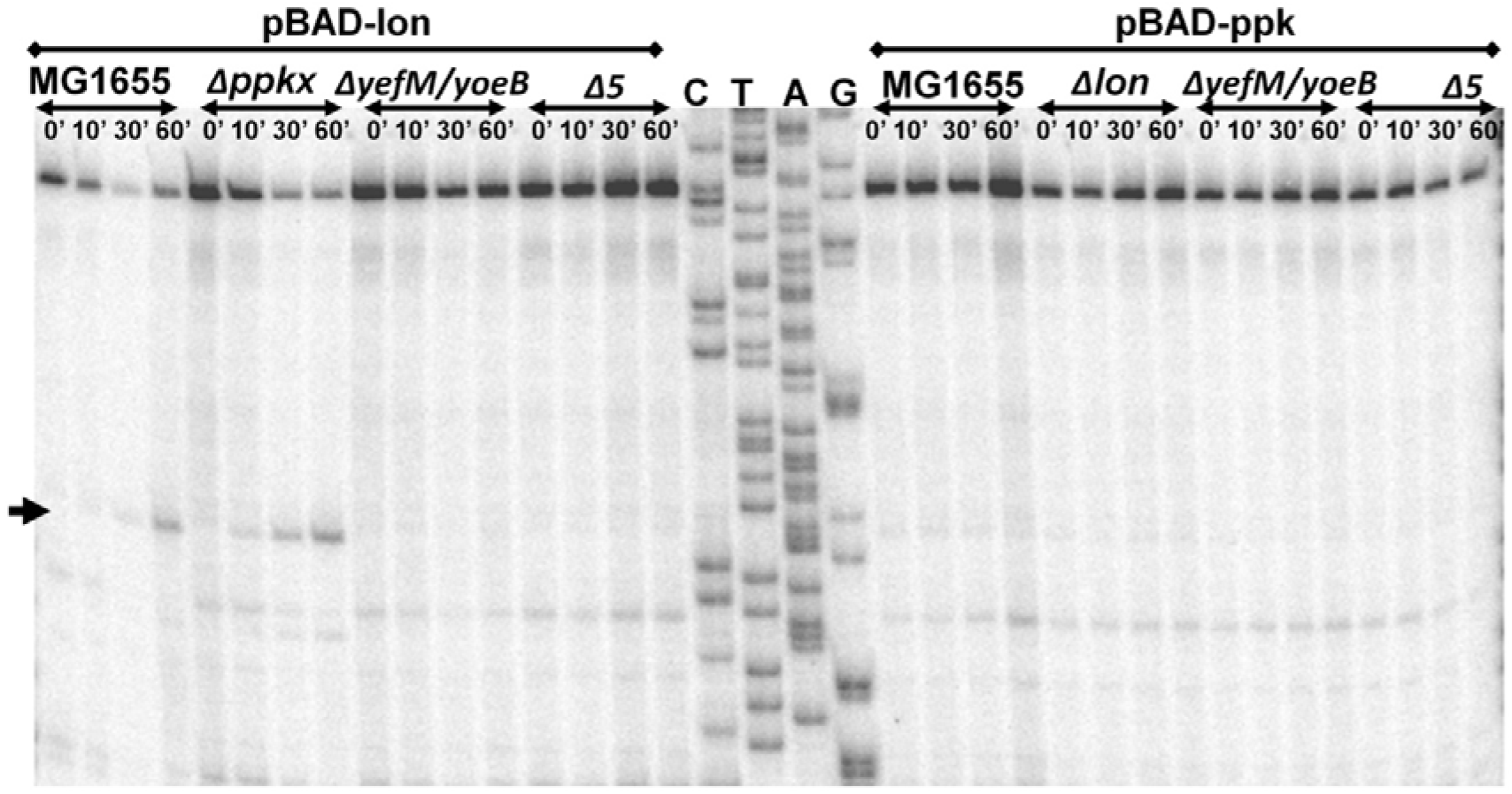
YoeB-dependent cleavage upon overexpression of *lon* is independent of polyP. MG1655, Δ*ppk*Δ*ppx (*Δ*ppkx)*, Δ*yefM/yoeB* and Δ*5* strains were transformed with pBAD-*lon* and pBAD-*ppk* was transformed into MG1655, Δ*lon*, Δ*yefM/yoeB* and Δ*5*. The transformants were grown in LB media, supplemented with 2% glycerol, to mid-exponential phase (0.45 of OD_450_) and 0.2% arabinose was added to induce expression of *lon* or *ppk*. Samples were collected at 0, 10, 30 and 60 minutes and primer extension was carried out using Lpp mRNA-specific primer (lpp21) for cleavage site mapping. YoeB-dependent cleavage, indicated by an arrow, is in accordance with results from Christensen, *et al*, 2004.

### Increase in the rate of transcription of *yefM/yoeB* loci upon heat shock

Several descriptions of TA regulations assumed TA loci to be bistable, either ON or OFF. This notion is strengthened by experiments in which starvation is drastic and near absolute which results in increased rates of TA transcription (Christensen, et al. 2001, Maisonneuve, et al. 2011). We hypothesized that TA regulation is rather “analogue” and not “discrete”, meaning that the degree of repression varies as a function of global translation and/or proteolysis rates. We performed heat shock experiments and oxygen deprivation experiments to mimic suboptimal conditions which could affect translation rates. Heat shock experiments were performed from 30°C to 42°C and 37°C to 47°C. In the 30°C to 42°C heat shock experiment, we did not notice any significant difference in the transcriptional rates of TAs. However in the 37°C to 47°C heat shock experiment, we noticed a stable threefold increase in the rate of transcription which was maintained through the 120 minutes of analysis (Figure 4). We performed an experiment in which cultures were deprived of oxygen by placing the cultures at 37°C in the incubator without shaking. Similar to the results in the 37°C to 47°C heat shock experiment, we observed that the *yefM/yoeB* transcript is consistently upregulated upon partial anaerobiosis (Figure 4). In fact, similar stable maintenance of increased transcription was also observed in *relBE* system upon induction of heat shock and glucose starvation (Christensen, et al. 2001).

**Figure 4.**
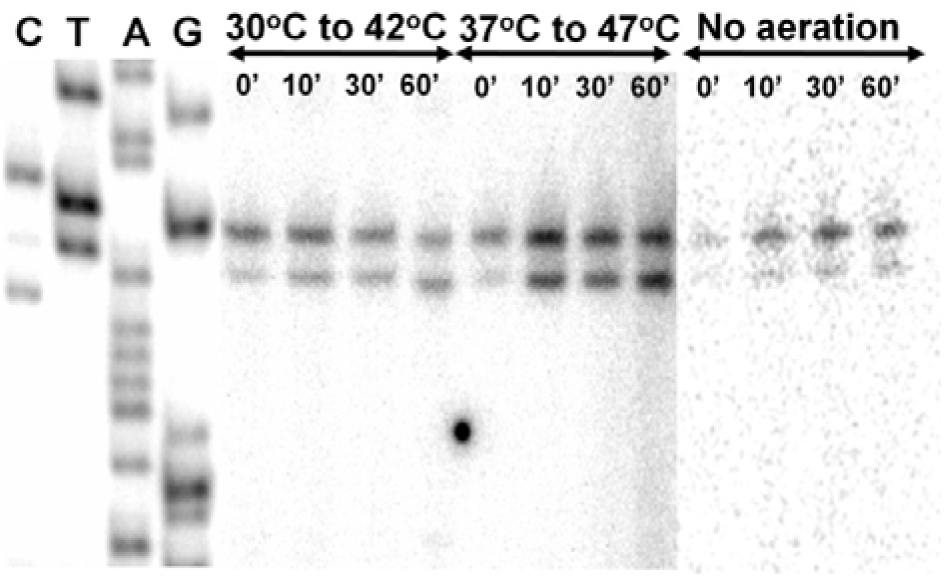
Effect of heat shock on the transcription of *yefM/yoeB* loci. Exponentially growing MG1655 cultures (0.45 of OD_450_) were moved from 30°C to 42°C, or 37°C to 47°C or aerated to unaerated conditions. Samples for RNA purification were collected at time 0, 10, 30 and 60 minutes after stress induction. Primer extension was performed on 10μg of each RNA sample using YefM mRNA-specific primer (YefMPE-2). The experiment labelled “no aeration” was conducted independently at a different time.

**Figure 5.**
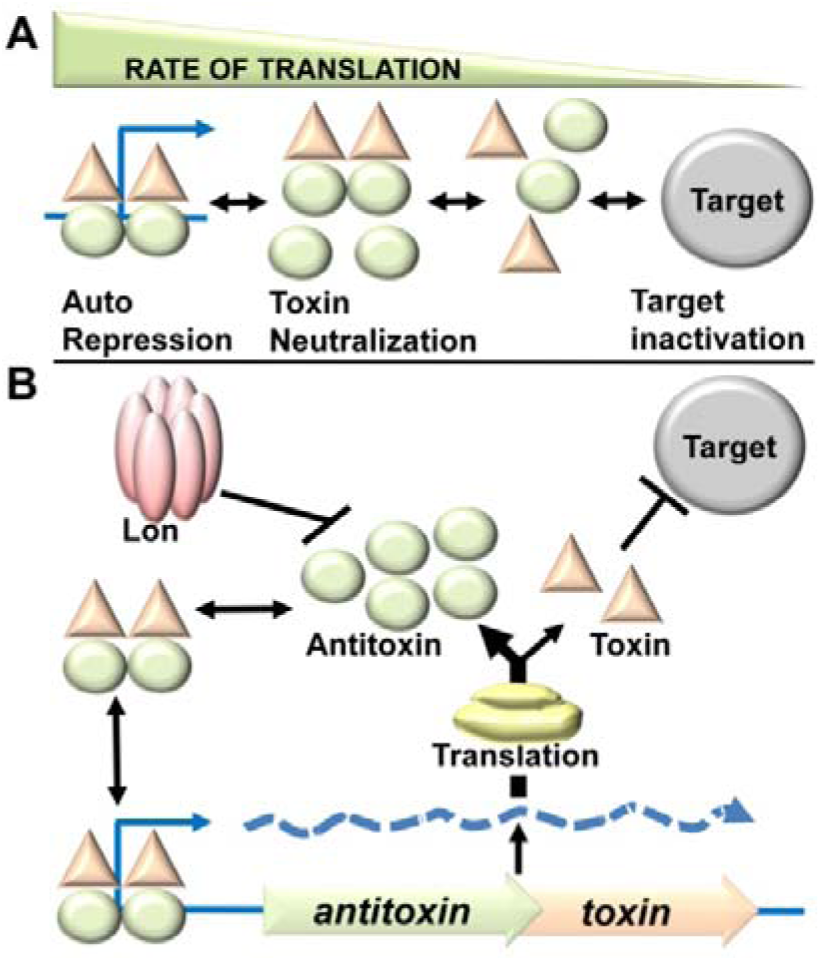
(**A**). **Interactions and action of toxin antitoxin proteins**. Based on the concentrations of antitoxin and toxin proteins the equilibrium shuttles between auto repression or target inactivation. The concentration of TA proteins is a function of rate of translation, assuming constant or negligible variation in the rate of antitoxin proteolysis by Lon. At very high translation rates the formation of repressive TA complexes results in repression of TA transcription. As the translation rate reduces the equilibrium shifts towards target inactivation. (**B**) **The dymanic regulatory circuit of Type II TA systems**. In all conditions, the rate of degradation is constant or negligibly variable. At higher translation rates more antitoxin proteins are produced per TA mRNA. The formed TA proteins form a complex in conditions of high translation rate which binds to the operator of TA operons and represses it.

## DISCUSSION

### PolyP is not essential for the transcriptional activation of *yefM/yoeB* loci and endoribonuclease activity of YoeB

In our amino acid starvation experiments, we found a consistent increase in transcription of *yefM/yoeB* operon in Δ*ppk*Δ*ppx* and Δ*relA*Δ*spoT* strains, strains deficient in accumulation of polyP and ppGpp respectively, similar to that of wild type (Fig 1A). In contrast to the reports of Maisonneuve et al., 2013, our semi-quantitative primer extension experiments show that neither ppGpp nor polyP is required for the transcriptional upregulation of *yefM/yoeB* loci. This indirectly indicates that ppGpp or polyP is not required for YefM degradation during amino acid starvation. Our results corroborate the earlier findings that *relBE* (Christensen, et al. 2001) and *mazEF* systems (Christensen, et al. 2003) transcriptional upregulation of during SHX-induced starvation is independent of ppGpp but dependent on Lon protease. RelA dependent accumulation of ppGpp was shown to be inhibited by chloramphenicol treatment (Svitil, et al. 1993) and yet the *relBE* and *mazEF* TAs were shown to be upregulated upon addition of chloramphenicol (Christensen, et al. 2001, Christensen, et al. 2003). When we overexpressed Lon protease in MG1655 and Δ*ppk*Δ*ppx*, there was a dramatic and consistent increase in transcription from *yefM/yoeB* loci (Fig 1C) in both the strains indicating that polyP is not required for degradation of YefM. In our overexpression experiments, we found a transient increase in transcription from *yefM/yoeB* operon upon overproduction of *relA* (Figure 2B) and *ppk* (Figure 2C) which could be due to metabolic burden of over-producing proteins. There is also a possibility that ppGpp production and/or polyP accumulation could affect the translational apparatus as in polyP-dependent Lon-mediated proteolysis (Kuroda, et al. 2001) of ribosomal proteins and hence the transient increase in *yefM/yoeB* transcription was observed. Previously, Lpp and tmRNA were shown to be cleaved by chromosomally encoded YoeB upon overproduction of Lon protease (Christensen, et al. 2004). We performed similar experiments to test the role of polyP in manifestation of YoeB-dependent endoribonuclease activity. We observed YoeB-dependent cleavage of Lpp mRNA, upon Lon overexpression, in MG1655 as well as in Δ*ppk*Δ*ppx* strains but not in Δ*yefM/yoeB* and Δ*5* (Fig 2). This implies that YoeB-specific cleavage is independent of polyP—meaning that activation of YoeB, by degradation of YefM, is independent of polyP. Furthermore, overexpression of *ppk* did not induce any YoeB-mediated cleavage in any of the strains. Hence, our results establish that polyP is not required for the transcriptional activation of *yefM/yoeB* loci and endoribonuclease activity of YoeB which imply that polyP is not required for Lon-mediated degradation of YefM. Within the scope of the experiments it can be argued that translation and Lon protease are the regulators of YefM concentration.

### Is polyP required for antitoxin degradation?

Reduction in persisters was also observed upon *relA* overexpression in Δ*10* and Δ*ppk*Δ*ppx* strains prompting Maisonneuve et al., 2013 to assume that degradation of the other antitoxins in *E. coli* (ChpS, DinJ, MazE, MqsA, HicB, PrlF, YafN, HigA) was also dependent on polyP (Maisonneuve, et al. 2013). It is known that most of the antitoxins are loosely folded or natively unfolded (Cherny and Gazit 2004, Nieto, et al. 2007) and is probably the reason for high turnover rates of antitoxins. The half-life of free native YefM in *E. coli* MC4100 strain was about 48 minutes (Cherny, et al. 2005) while that of his-tagged YefM, in MG1655, is about 11 minutes (Maisonneuve, et al. 2013). The difference in these values could be attributed to differences in the genetic background of the strains as well as artificially induced primary structural modifications in YefM. It is to be noted that YefM is degraded (Cherny, et al. 2005) even in MC4100 strain (*relA1* mutant strain) which is deficient in accumulating ppGpp during amino acid starvation (Metzger, et al. 1989). It may be noted that antitoxins like YafN, HigA and YgiT were shown to be degraded by both Lon and Clp proteases (Christensen-Dalsgaard, et al. 2010). Furthermore, based on studies on “delayed relaxed response” (Christensen and Gerdes 2004), the half-life of RelB in MC1000 strain is approximately 15 minutes and RelB101 (A39T mutant of RelB) is less than 5 minutes. It is interesting to notice that RelB101 is degraded even in a Δ*lon* strain, indicating that some other proteases may also cleave RelB101 (Christensen and Gerdes 2004). This literature evidence indicates that changes in primary structures of antitoxins could drastically alter their stabilities and protease susceptibility. Maisonneuve et al., 2013 did not provide accurate half-life of YefM in Δ*ppk/*Δ*ppx* strain (Maisonneuve, et al. 2013) to ascertain if there is any degradation at all. The “polyP-dependent active TA regulation model” (Maisonneuve, et al. 2013) fails to explain how all the ten significantly divergent antitoxins of *E. coli* MG1655 could be substrates of ‘polyP-modulated Lon’ protease. The requirement of polyP for Lon-mediated degradation of antitoxins does not seem to be a general phenomenon unless ‘native unfoldedness’ and/or nucleic acid binding motif (the two important features of antitoxins) of the substrate protein determine it. Moreover, the molecular mechanisms of Lon substrate specificity modulation by polyP are not yet fully understood. It is not yet clear if a specific motif or tertiary structure of the substrate protein determines polyP-Lon dependent proteolysis. PolyP was shown to inhibit Lon protease *in vitro* (Osbourne, et al. 2014) and is implicated in acting as a chaperone for unfolded proteins (Kampinga 2014) which may have significant implications in bacterial stress physiology. To our rationale, since TAs propagate through horizontal gene transfer mechanisms, minimal dependence on host genetic elements maybe preferable for regulation. Within the scope of our experiments conducted in this study and based on evidence in literature, it is appropriate to state that polyP is not essential for Lon-mediated proteolysis of YefM. Although we do not have a ready explanation for this fundamental contradiction, we do not rule out His-tag interference in the proteolysis assays used by Maisonneuve et al., 2013.

### Dynamic model of TAs

Our experiments (Figure 2, 3) rule out the essentiality of ppGpp and polyP in the transcriptional regulation of *yefM/yoeB* loci and also in the active model. Our results largely reiterate the so called “passive model” (Gerdes, et al. 2005, Sat, et al. 2001) which we refer to as “dynamic model”. In the dynamic model, production by translation and continuous degradation by protease contribute to high turnover rates of antitoxins. It is assumed that degradation by protease is relatively constant and the production of antitoxin is highly variable according to the growth conditions that influence translation. Regulation of TAs is not just the discrete ‘ON’ or ‘OFF’ states at the transcriptional level but rather a function of the continuous variable “antitoxin concentration”. Same bacteria in different conditions will have different levels of antitoxin which could reflect in the transcriptional up/down regulation of the corresponding TA operon. This could be a sensory mechanism for the TAs to detect the nature of the growth conditions and thus modulate its regulation. The key to the dynamicity of the TA regulatory system is the continuous degradation of antitoxin likely by virtue of the loose conformation of the antitoxin, a characteristic feature of most Type II antitoxins. Since translational inhibition precedes proteolysis during conditions like amino acid starvation, it would be rational for TAs to be responsive to translational inhibition rather than the proteolysis of antitoxin. Towards establishing non-discrete nature of the TA operon regulation we performed heat shock and oxygen deprivation experiments (Figure 4).

Interestingly, we found that at 47 °C and O_2_ deprivation there is a consistent threefold increase in the transcription rate of *yefM/yoeB* loci indicating that the regulation of *yefM/yoeB* is not just an ON or OFF but has intermediate states. Heat shock response alleviates the effects and burden of non-productive proteome of the cell by production of more chaperones and proteases including Lon. Although it is beyond the scope of the experiment to pinpoint the cause, it is likely that the increased rate of transcription is due to higher turnover rate of YefM due to structural changes in the cell. Experiments involving drastic and instantaneous inhibition of translation, like treatment with high doses of SHX or Chloramphenicol, the antitoxins are almost completely degraded and hence we observe a dramatic increase in transcription over time. As opposed to starvation experiments in which TA mRNA seems to accumulate over time, stable maintenance of transcriptional upregulation indicates that the operator is partially or intermittently repressed. In fact, similar stable maintenance of increased transcription was also observed in *relBE* system upon induction of heat shock and glucose starvation (Christensen, et al. 2001). Stresses like heat shock, anaerobic conditions and glucose starvation reduce productive translation which could reflect in lower rates of antitoxin production. This leads to lesser probability for the formation of TA complexes capable of autorepression. Hence an increased transcription rate is maintained from the TA loci and is a function of the new equilibrium of translation and proteolysis rates of antitoxin. This dynamic nature of TA regulation allows sensing the global translation rate which usually indicates the nature of the growth conditions and available nutrient resources.

Interestingly, in the abstract of a recent report (Germain, et al. 2015) a statement was made; “Polyphosphate activated Lon to degrade all known type II antitoxins of *E. coli*.”. Firstly, there is no evidence ever provided that ‘all known antitoxins’ were degraded by polyP modulated Lon in any report and moreover such observation is questionable. The results presented in our report and the exhaustive literature survey conclusively refute the essentiality of ppGpp and polyP in the regulation of *yefM/yoeB* and likely other similarly working TAs.

## MATERIAL AND METHODS

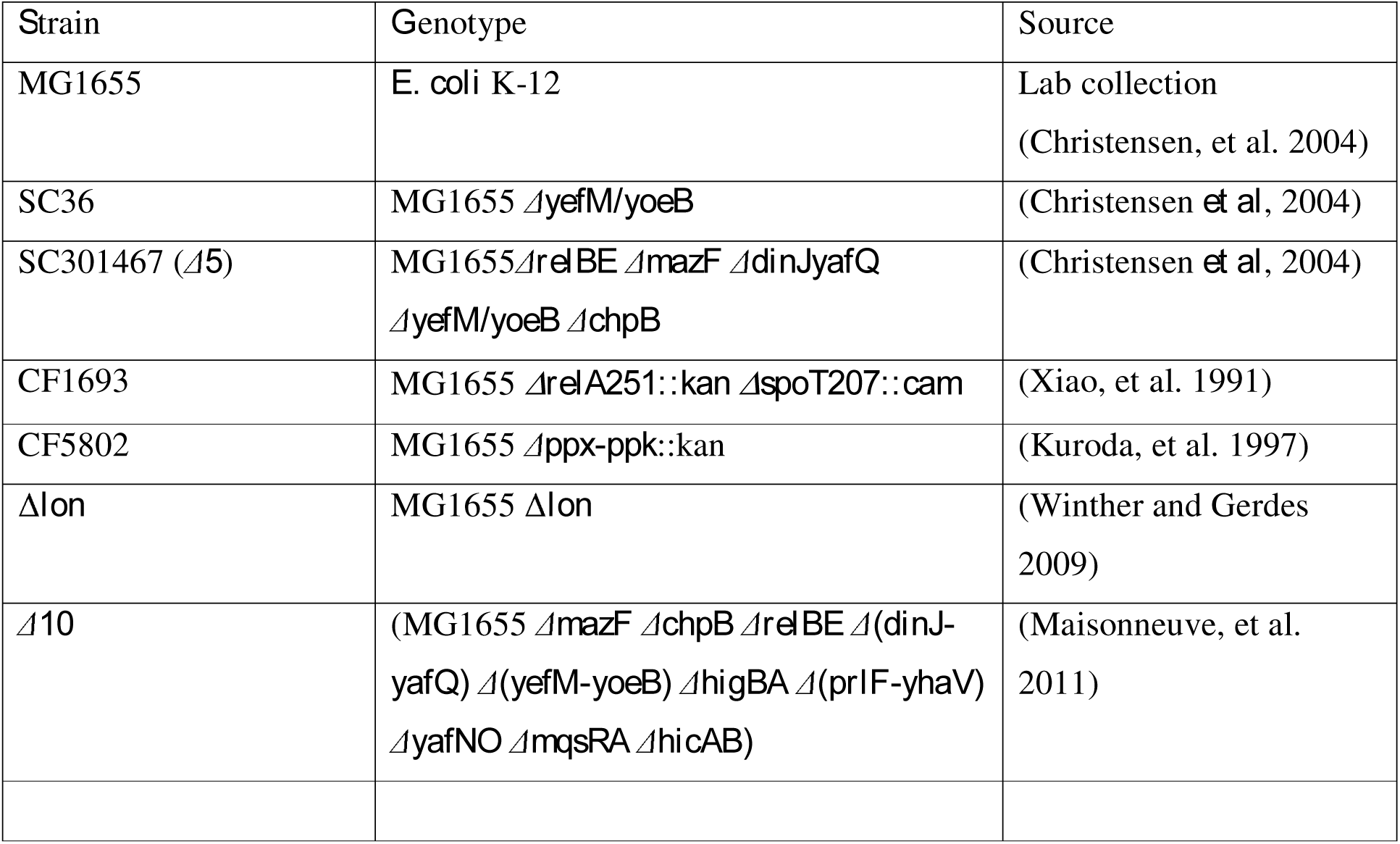

| Plasmid |  | Source |
| --- | --- | --- |
| pBAD33 |  | (Guzman, et al. 1995) |
| pBAD-*lon* |  | (Christensen, et al. 2004) |
| pBAD-*ppk* |  | Mikkel Girke Jorgensen,<br>unpublished |
| pALS13 | AmpR/ Ptac -*relA'* (RelA 1-455), active<br>RelA | (Svitil, et al. 1993) |
| pUC18 |  |  |

### Growth conditions and media used

All the experiments involving primer extension were grown in Luria Bertani broth, at 37 °C, with 180 rpm shaking in a water bath unless specified otherwise. Primer extension Samples of 25 mL experimental cultures were collected at 0, 10, 30 and 60 minutes and cells were harvested by centrifugation at 4°C. Total RNA was isolated using hot phenol method and quality was analyzed by agarose gel electrophoresis. Total RNA in each sample was set to about 5 μg/μL. p32 labelled primers, YefMPE-2 (5’-GGCTTTCATCATTGTTGCCG-3’) and lpp21 (5’-CTGAACGTCAGAAGACAGCTGATCG-3’), were used in primer extension experiments involving *yefM/yoeB* promoter activity and YoeB-dependent mRNA cleavage site mapping respectively. Reverse transcription was carried out on 10 μg of total RNA, purified from samples at designated time points, using AMV-reverse transcriptase. Sequencing reactions were carried out similarly with Sanger’s dideoxynucleotide method.

Antibiotic sensitivity assay:

Conventional disc diffusion method was used to measure the relative sensitivity of the strains. 100 μL of diluted (100-fold) overnight cultures were spread on LB agar (height – 5 mm) in plates with diameter 9.5 cm. Premade antibiotic discs with defined concentrations (purchased from HiMedia^TM^) were placed on the agar plates after 20 minutes. The plates were incubated overnight at 37 °C. Diameters of the zones of inhibition were measured and the graph was plotted.

## ACKNOWLEDGEMENT

The author thanks Prof. Kenn Gerdes for the support during this work at BMB, University of Southern Denmark.

